# VZV gE Promotes STING Degradation via Midnolin-Proteasome Pathway to Inhibit cGAS-STING Signaling

**DOI:** 10.1101/2025.01.13.632693

**Authors:** Lumiao Chen, Lingao Dai, Shenghao Nie, Yue Zhao, Fanyuan Li, Xiujiao Deng, Chunlin Liu, Moran Li, Zewei Luo, Yuqing Liu, Weiwei Ge, Xiaofei Liu, Qian Yi, Yongkui Li, Pan Pan, Xiaoping Wang

**Affiliations:** Department of Pain Management, the First Affiliated Hospital of Jinan University, Guangzhou 510630, China; Key Laboratory of Viral Pathogenesis & Infection Prevention and Control (Jinan University), Ministry of Education, Guangzhou 510632, China, College of Life Science and Technology, Jinan University, Guangzhou 510632, China; The First Affiliated Hospital of Jinan University, Guangzhou 510630, China; MOE Key Laboratory of Laser Life Science & Institute of Laser Life Science, College of Biophotonics, School of Optoelectronic Science and Engineering, South China Normal University, Guangzhou 510631, China; School of Basic Medical Science, State Key Laboratory of Respiratory Disease, Guangzhou Medical University, Guangzhou 511436, China

**Author notes:** Corresponding author: Xiaoping Wang, Department of Pain Management, the First Affiliated Hospital of Jinan University, Guangzhou 510630, China. Co-corresponding authors: Yongkui Li,; Pan Pan.

**Keywords:** Varicella-zoster virus (VZV)| glycoprotein E (gE)| STING interaction| Antiviral immune responses| Proteasome pathway

## Abstract

Varicella-zoster virus (VZV) infects 85 million individuals annually and it is known for its intricate interplay with host cellular mechanisms, significantly impacting antiviral immune responses. Previous studies have highlighted the critical role of VZV glycoprotein E (gE) in virus-host interactions, but the precise mechanisms remain to be elucidated. Our results demonstrate that VZV gE interacts with STING, specifically through the gE (350-480aa) and STING (151-160aa) regions. This interaction predominantly involves incompletely glycosylated gE. Functional assays revealed that gE inhibits the cGAS-STING signaling pathway. gE enhances VZV proliferation and promotes STING degradation via the proteasome pathway, without affecting STING ubiquitination. We identified Midnolin as a mediator in this degradation process, with gE facilitating the interaction between STING and Midnolin. In vivo, gE expression in mice led to diminished antiviral responses upon HSV-1 infection, highlighting gE’s role in modulating immune signaling. Our findings provide significant insights into VZV’s evasion of host immune responses. By promoting STING degradation through an unconventional Midnolin-proteasome pathway, VZV gE effectively suppresses antiviral signaling, facilitating enhanced viral proliferation. The findings contribute significantly to the understanding of VZV pathogenesis and provide new therapeutic strategies against VZV.

## Introduction

VZV infections pose a significant global health burden, often presenting without prodromal symptoms before the characteristic rash emerges. This virus is renowned for causing chickenpox in children and adolescents, remaining latent for years or decades before potentially reactivating as shingles, often without a discernible trigger [1]. Despite the advent of the attenuated live vaccine for chickenpox in 1974 and a specific vaccine for shingles in 2006 [2] [3], their universal inclusion in national immunization programs remains an exception rather than the rule in many countries. In 2019, VZV infections led to a staggering 84 million cases worldwide and contributed to a significant loss of 900,000 Disability-Adjusted Life Years (DALYs) [4]. Moreover, the global pandemic of COVID-19 has exacerbated the incidence of herpes zoster (HZ) [5], further underscoring the persistent and severe nature of this viral menace.

Our understanding of the intricate infection processes and the mysterious reactivation of VZV from latency remains confined. This is compounded by the virus’s extensive genome, which encodes an impressive number of approximately 71 open reading frames (ORFs) [6]. Among the 10 glycoproteins on its envelope, glycoprotein E (gE) stands out as the most abundant and crucial for the virus’s proliferation cycle. The unique N-terminus of gE plays a dual role in viral proliferation and disease manifestation[7] [8] [9]. Previous studies have demonstrated that the 27-90 amino acid segment of VZV gE is capable of interacting with the insulin-degrading enzyme (IDE), thereby potentiating the virus’ infectivity [10] [11]. Moreover, the distinctive role of gE’s N-terminus in facilitating T-cell tropism is crucial for its neurovirulent characteristics [12]. Recently, the development of recombinant gE protein adjuvant for shingles prevention has marked a significant advancement in vaccine technology [13] [14]. Despite these advancements, the intricate interplay between VZV and the host immune system, particularly involving gE and the cGAS-STING signaling pathway, remains largely unexplored.

The cGAS-STING (cyclic GMP-AMP synthase-Stimulator of Interferon Genes) signaling cascade is crucial for the innate immune defense of the host. Upon stimulation by DNA viruses, this pathway becomes active, centered around STING, which forms a complex with TANK-binding kinase 1 (TBK1) and Interferon regulatory Factor 3 (IRF3) [15]. This complex then migrates to the perinuclear area, triggering a cascade of events leading to the expression of type I interferons and inflammation-related genes [16]. The endogenous STING protein undergoes rapid degradation post-DNA stimulation, making the cGAS-STING-induced interferon signaling transient and highly responsive. Research by Dobbs et al. revealed that STING degradation initiates during its transition from the endoplasmic reticulum to the Golgi apparatus [17], a process distinct from the ubiquitination pathway controlled by iRhom2 [18]. Gonugunta et al. further elucidated STING’s lifecycle, demonstrating that following interferon signaling, STING is directed to Rab7-positive endolysosomes for degradation [19]. This degradation process depends on the presence of the 281-297aa motif and can be blocked by bafilomycin A1. Notably, the rapid turnover of STING occurs independently of TBK1, IRF3, downstream signaling, ubiquitination, or S366 phosphorylation, underscoring the sophistication and complexity of the regulatory mechanisms governing this crucial immune signaling molecule.

Given the significance of the cGAS-STING pathway in innate immune defense, we investigated its interaction with VZV gE. Proteasomal degradation is a vital route for intracellular protein breakdown, including STING, with the proteasome—a formidable complex—skillfully recognizing and dismantling Ub-tagged proteins, primarily via K48-linked Ub chains [20] [21]. Recent research has sparked new interest in proteasomal studies by unveiling a unique, non-Ub-dependent degradation pathway regulated by midnolin [22]. Midnolin accelerates proteasomal degradation of substrate proteins, using its Ub-like domain to bypass traditional ubiquitination. Various transcription factors from the immediate early gene (IEG) family, transiently activated by extracellular signals, undergo swift degradation via the midnolin-proteasome pathway. However, whether STING might also be degraded through this pathway remains unclear, necessitating further exploration of this proteasomal intricacy.

Our study uncovers a novel mechanism by which the under-glycosylated form of gE interacts with STING, enhancing its degradation through the midnolin-proteasome pathway, thereby modulating the host immune response. Consistent with this, a murine model exhibited increased pathological damage upon HSV1 infection in the presence of gE. Furthermore, our research revealed a novel mechanism for STING degradation mediated by the midnolin-proteasome pathway, which gE was found to enhance. This finding offers fresh insights into the interplay between VZV and its host, emphasizing gE’s role in manipulating the host immune response. Understanding the connection between gE and the cGAS-STING pathway is not merely academic; it holds promise for vaccine and therapeutic development against viral infections. By deciphering these molecular interactions, we are paving the way for more effective strategies to combat VZV and other viral threats.

## Results

### VZV gE interacts with STING

VZV glycoprotein synthesis and trafficking are similar to HSV1, with glycoproteins moving from the endoplasmic reticulum to the Golgi and then to the cell surface, before being internalized to the trans-Golgi network for virion envelopment [23]. In an effort to delineate the interactions between VZV glycoproteins and the STING, this investigation utilized a VZV-GFP construct sourced from the Dumas strain as a reference. Based on this, plasmids encoding 10 distinct glycoproteins, each appended with a FLAG epitope tag for detection, were engineered. Co-IP results revealed several glycoproteins interacted with HA-STING, included gB, gM, and gE (Figure 1A). As the preeminent glycoprotein expressed by VZV and a pivotal target for vaccine development, the engagement of gE with STING is compelling and demands deeper inquiry. Further results also confirmed the interaction between STING and gE (Figure 1B).

**Figure 1.**
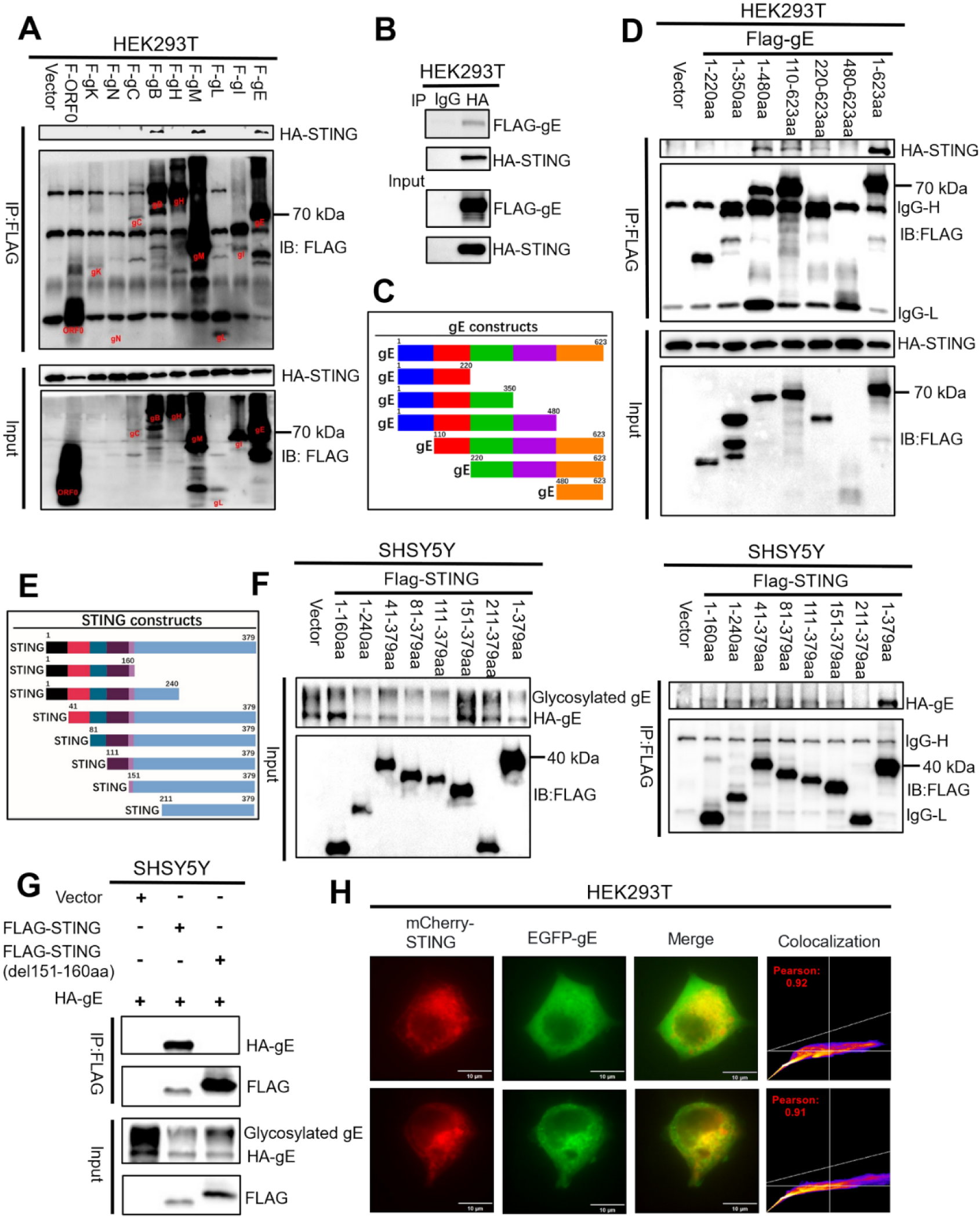
VZV gE interacts with STING. A: VZV Glycoprotein-STING Interactions: A series of individual FLAG-tagged VZV glycoproteins were co-transfected with HA-STING into 293T cells. Samples were harvested at 24 hours post-transfection for subsequent analysis. B: FLAG-gE and HA-STING Co-transfection Assay: FLAG-gE was co-transfected with HA-STING in 293T cells, and samples were collected 24 hours later for further testing. C: Schematic Representation of FLAG-gE Truncated Plasmids. D: Analysis of FLAG-gE Truncated Plasmids’ Interaction with HA-STING: A variety of FLAG-tagged gE truncated plasmids were co-transfected with HA-STING into 293T cells, followed by sample collection 24 hours later for detailed examination. E: Schematic Representation of FLAG-STING Truncated Plasmids. F: FLAG-STING Truncated Plasmids’ Interaction with HA-gE: Diverse FLAG-tagged STING truncated plasmids were co-transfected with HA-gE into SHSY5Y cells, with samples obtained 24 hours post-transfection for analysis. G: FLAG-STING and FLAG-STING (del151-160aa) plasmids’ Interaction with HA-gE: plasmids were co-transfected with HA-gE into SHSY5Y cells, with samples obtained 24 hours post-transfection for analysis. H: Confocal Microscopy of EGFP-gE and mCherry-STING Co-transfection: EGFP-gE (green) and mCherry-STING (red) were co-transfected into 35 mm confocal dishes to facilitate imaging using confocal microscopy.

In pursuit of elucidating the precise domain within gE responsible for this interaction, plasmids harboring the wild-type (WT) gE sequence alongside six distinct truncated variants were designed (Figure 1C). Among these, the truncated forms gE (1- 480aa), gE (110-623aa), and gE (220-623aa) demonstrated interaction with STING (Figure 1D), suggesting that gE (350-480aa) is required for the interaction with HA- STING. Similarly, plasmids harboring the full-length STING sequence and its seven truncated versions were introduced to explore its binding area with gE (Figure 1E). Results unveiled that the STING fragments STING (1-160aa), STING (1-240aa), STING (41-379aa), and STING (81-379aa), STING (111-379aa), and STING (151-379aa) were in intimate conversation with HA-gE (Figure 1F). Additional findings revealed that the STING protein lacking the 151-160aa sequence fails to interact with HA-gE, indicating that this sequence is essential for binding to HA-gE (Figure 1G).

Confocal microscopy studies confirmed a significant colocalization between mCherry-STING and EGFP-gE, indicating a strong interplay, and underscored by robust Pearson coefficients reaching 0.92 and 0.91, respectively (Figure 1H).

The above results demonstrated that the interaction between gE (350-480aa) and STING (151-160aa), revealing a potential regulatory link between VZV glycoproteins and STING’s functionality.

### The interaction with STING occurs with incompletely glycosylated VZV gE

In order to determine the VZV gE interacts with STING during VZV infection, we infected ARPE-19 or SHSY5Y cells with VZV-GFP. Amazingly, CO-IP assays proved that no interaction between gE and endogenous STING was observed at 24 or 48 hours post-infection in either ARPE-19 (Figure 2A-2C) or SHSY5Y cell lines (Figure 2D). It has been reported that the commercially available antibody targeting VZV gE only binds to partial conformational epitopes of VZV gE [11], which aligns with our findings (Figure 2E) showing that only the 110-350 amino acid region of FLAG-tagged gE can be recognized by the gE antibody sourced from Abcam (ab272686). This observation may explain why no interaction between STING and gE was detected during VZV-GFP infection.

**Figure 2.**
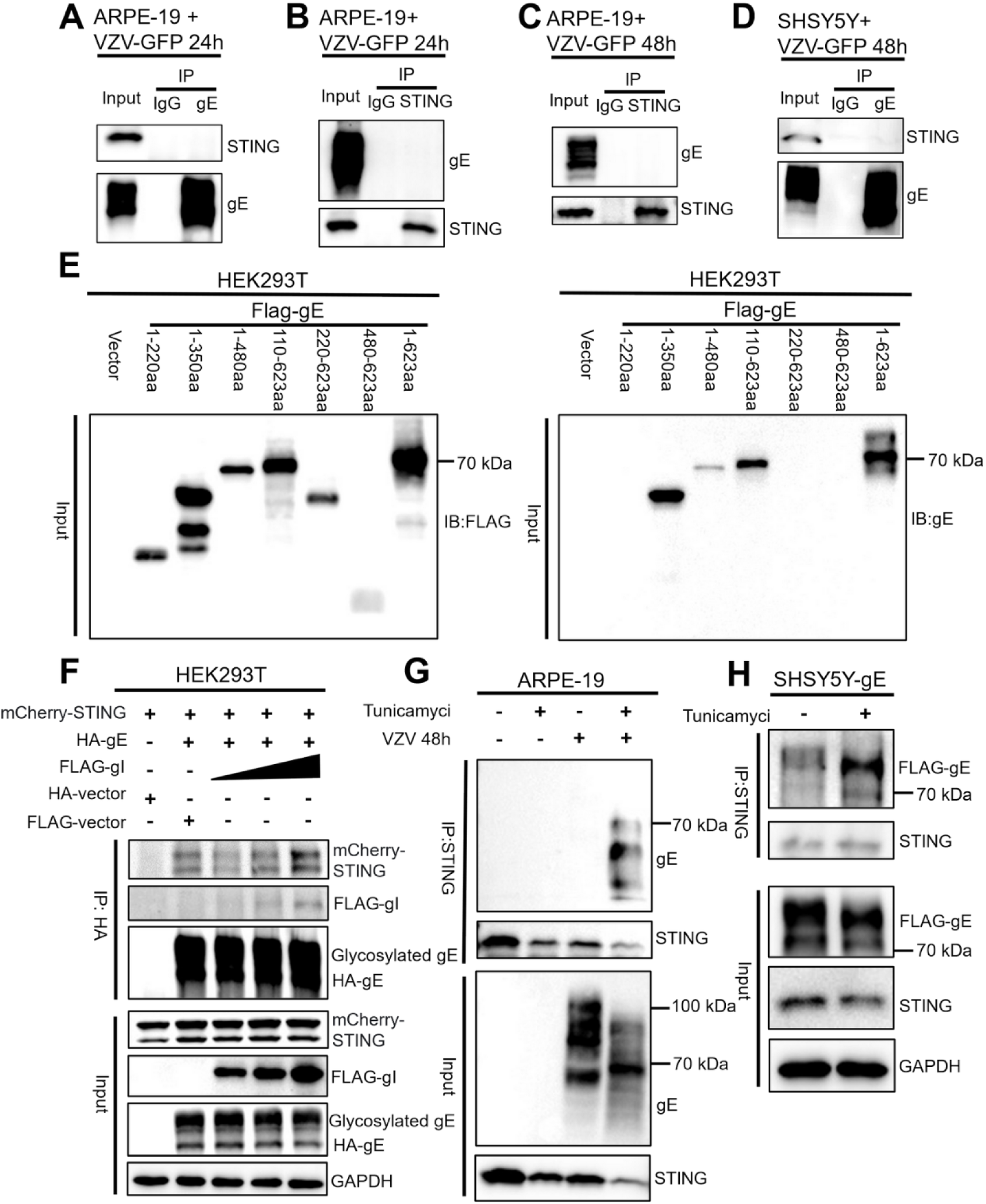
The interaction with STING occurs with incompletely glycosylated VZV gE. A, B, C, and D: Conduct endogenous gE/STING co-immunoprecipitation (CO-IP) experiments on cells infected with VZV-GFP. The infection conditions are as follows: ARPE-19 cells infected for 24 hours; ARPE-19 cells infected for 24 hours; ARPE-19 cells infected for 48 hours; SHSY5Y cells infected for 48 hours. E: WB results of FLAG-gE Truncated Plasmids, which were detected by FLAG antibody and gE antibody respectively. F : Results of the CO-IP experiment in 293T cells co-transfected with FLAG-gI, STING and with a concentration gradient along with gE. G: Results of the CO-IP experiment between STING and gE in ARPE-19 cells infected with VZV-GFP without treatment or with tunicamycin treatment (2 μg/mL for 24 hours); the cells were infected for 48 hours. H: Results of the CO-IP experiment between STING and gE in SHSY5Y-gE cells without treatment or under tunicamycin treatment (2 μg/mL for 12 hours).

Previous research suggests that gE and glycoprotein I (gI) form a heterodimer during VZV infection, enhancing the virus’s affinity for skin cells [24]. We then investigated whether this complex affects gE-STING interactions. Despite increasing gI concentrations, we found no change in the interaction between mCherry-STING and HA-gE, indicating that gE-STING interaction is not influenced by gI (Figure 2F).

The liaison between gE and IDE (Insulin Degrading Enzyme) transpires in the context of its immature, under-glycosylated state, identified as a 73-kDa species [11]. UniProt search indicates that the gE protein may possess several post-translational modification sites at its N-terminus, such as N-glycosylation at amino acid positions 266 and 437(Uniprotkb:P09259). Notably, the interaction domain of gE with STING includes the glycosylation site at 437aa (Figure 1D). Further experiments using tunicamycin, which inhibits glycosylation [25], showed an interaction between gE and STING in VZV-GFP-infected ARPE-19 cells (Figure 2G). The gE in this interaction was under-glycosylated, with a reduced molecular weight, indicating that gE in its immature form is the primary mediator of the interaction with STING during VZV infection. When we applied tunicamycin to SHSY5Y cells stably expressing gE, we also observed increased binding of gE to STING (Figure 2H). The above results demonstrated that the glycosylation level of gE could affect its interaction with STING during VZV infection.

### VZV gE inhibited cGAS-STING signaling pathway by targeting STING

To study gE’s impact on the cGAS-STING pathway, we used a dual-luciferase reporter gene assay in HEK293T cells, focusing on IFN-β promoter level. We found that gE significantly suppressed IFN-β promoter level, and this suppression was time-dependent (Figure 3A). The result also illuminated that a marked reduction in the levels of phosphorylated TBK1 (p-TBK1) protein within the U251-gE cells after VZV-GFP infection (Figure 3B). Extending our investigation to the impact of HSV-1 infection (the activator of cGAS-STING pathway), we observed a consistent theme of suppression; at 6, 9, and 12 hours post-infection, the U251-gE cells exhibited notably diminished mRNA levels of key inflammatory mediators, including IFN-β and TNF-α (Figure 3C).

**Figure 3.**
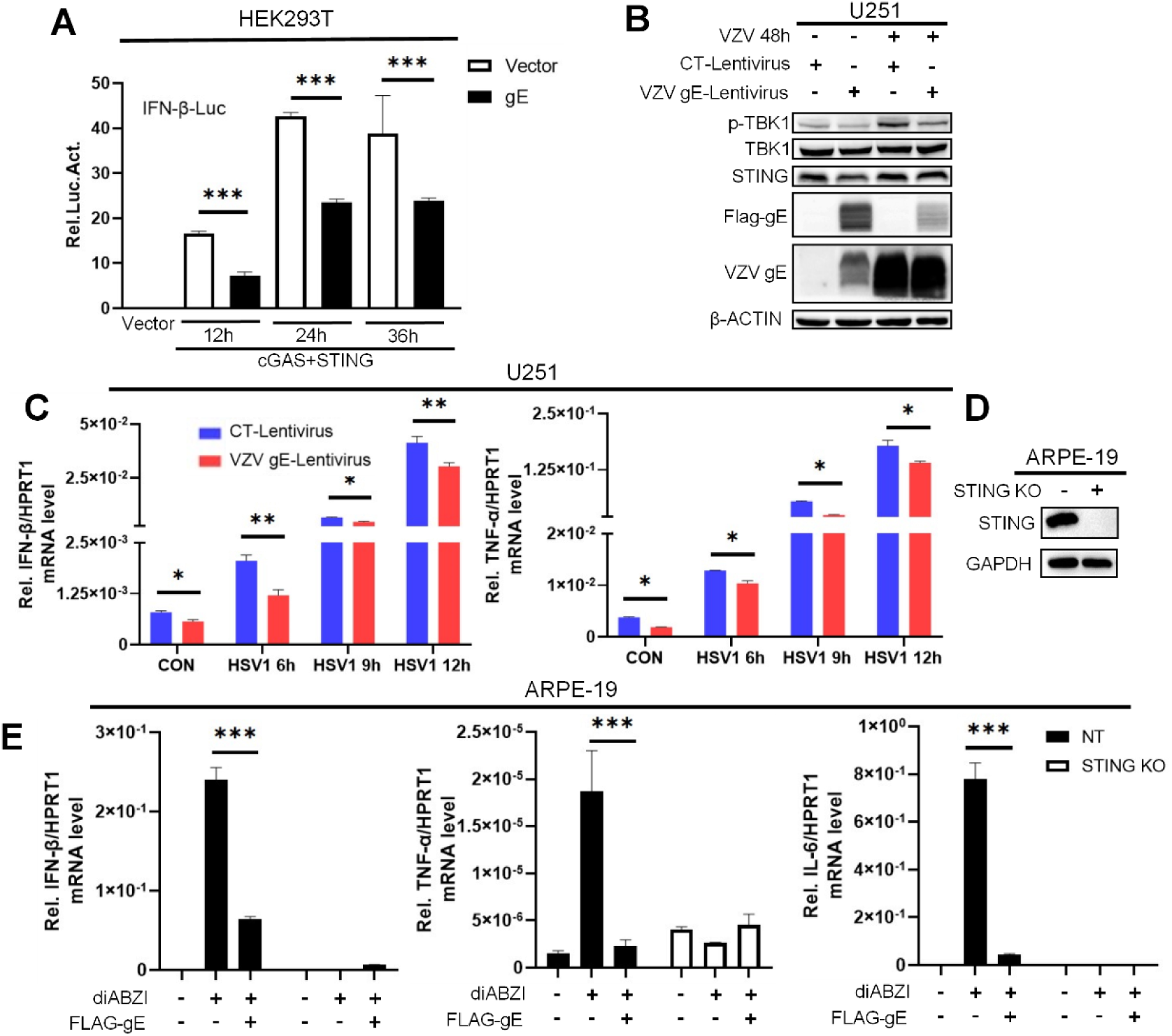
VZV gE targets at STING level to inhibiting cGAS-STING-triggered signaling pathway. A: The changes in IFN-β-Luc/TK results downstream of cGAS+STING at co-transfection times of 12, 24, and 36 hours under conditions without and with gE expression. B: The results of protein immunoblotting in U251-CT and U251-gE cells infected with VZV-GFP for 48 hours. C: The changes in mRNA levels of IFN-βand TNF-α in ARPE-19-CT/ARPE-19-gE cells without and with HSV-1 infection at 6, 9, and 12 hours. D: The WB results in ARPE-19 cells under NT (non-targeting) and STING knockout conditions. E : The changes in relative mRNA levels of IFN-β, TNF-α, and IL-6 in ARPE-19 STING NT and KO cells stimulated without or with diABZI treatment (5 μM, 1 hour).

Using CRISPR-Cas9, we knocked out the STING gene in ARPE-19 cells (Figure 3D). With diABZI, a STING activator, we found that the inhibitory impact of gE on inflammation-related genes (IFN-β, TNF-α and IL-6) was absent in STING-deficient cells (Figure 3E), suggesting that gE’s modulation of the cGAS-STING pathway is STING-dependent.

### gE enhances the proliferation of VZV-GFP

The VZV-GFP is derived form WT Oka strain, which incorporates a green fluorescent protein (GFP) and a luciferase gene into its genome, has demonstrated growth characteristics equivalent to the original Oka strain [26]. In studies with VZV-GFP infection in U251-CT and U251-gE cells, compared with U251-CT cells, U251- gE cells showed higher GFP expression and viral proliferation level (Figure 4A), suggesting gE enhances VZV proliferation . In SHSY5Y-CT and SHSY5Y-gE cells in conditions of T [27] unicamycin infecting with VZV-GFP, SHSY5Y-gE cells also had higher GFP expression, indicating increased viral proliferation (Figure 4B and 4C). Tunicamycin treatment, which affecting gE glycosylation and its interaction with STING, resulting in increased VZV fluorescent cluster count. Overall, the presence of gE appears to be beneficial for the proliferation of VZV, offering a nuanced perspective on the interplay between viral proliferation.

**Figure 4.**
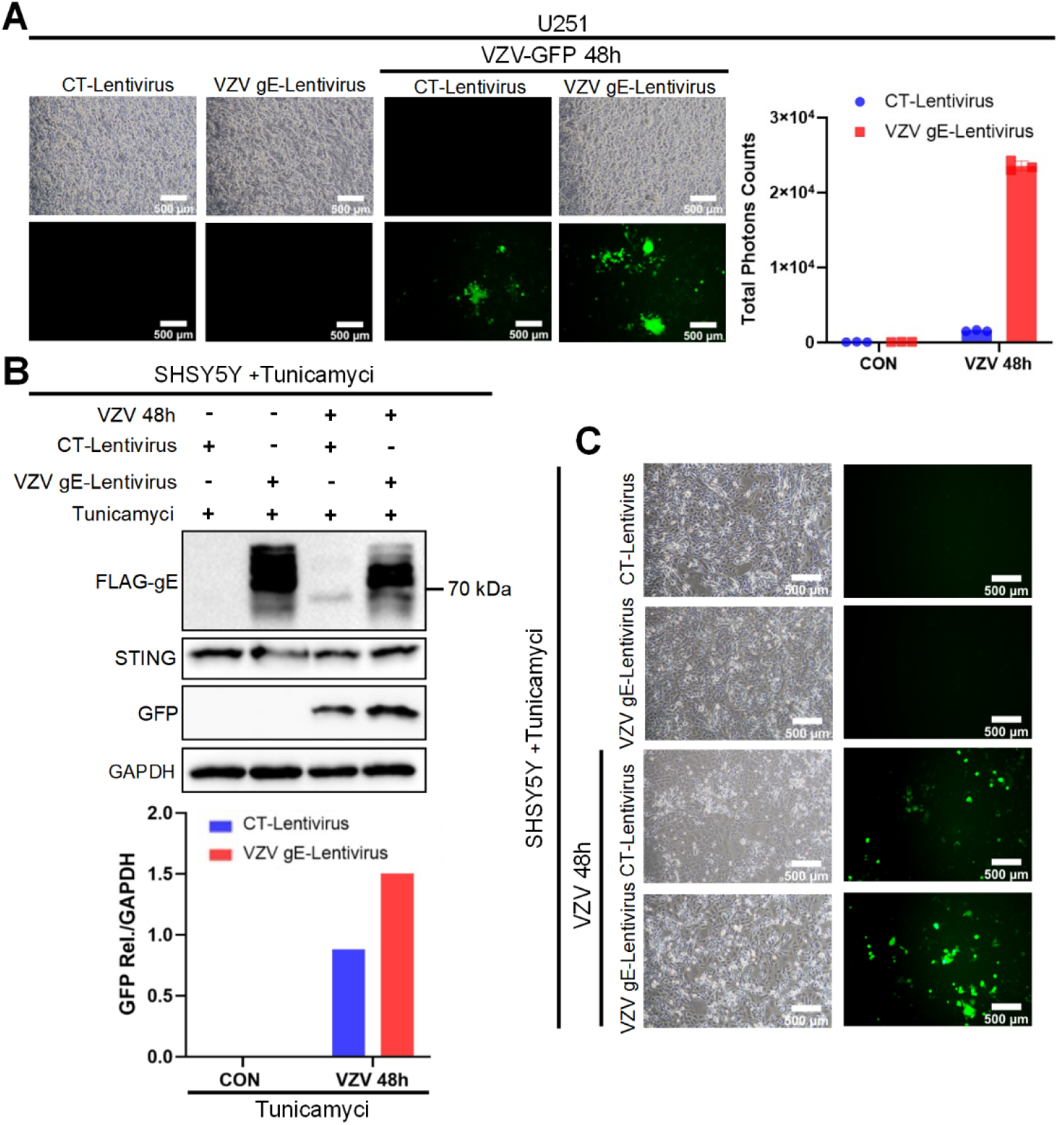
gE enhances the proliferation of VZV-GFP. A: Microscopic and fluorescence results of U251-CT and U251-gE cells infected with VZV-GFP for 48 hours. B: Protein immunoblotting results and GFP relative densitometry data for SHSY5Y-CT and SHSY5Y-gE cells infected with VZV-GFP for 48 hours. C: The fluorescent imaging results for SHSY5Y-CT and SHSY5Y-gE cells after a 48-hour exposure to VZV-GFP. Tunicamycin treatment cindition in B and C: 2 μg/mL for 24 hours.

### gE promotes the proteasome pathway to degrade STING

The cGAS-STING signaling pathway, once activated, initiates a cascade of protein degradation events that are exquisitely controlled. This regulatory finesse is paramount, ensuring a well-timed and calibrated immune response to foreign pathogens, while simultaneously averting the specter of unwarranted inflammation and the risks of autoimmune disorders [16]. In our experiments, the activation of this pathway by HSV- 1 led to a decrease in STING protein levels, hinting at a regulatory interplay (Figure 5A), which is consistent with the literature reports [27]. Additionally, upon stimulation with diZBAI, STING was also noticeably degraded (Figure 5B).

**Figure 5.**
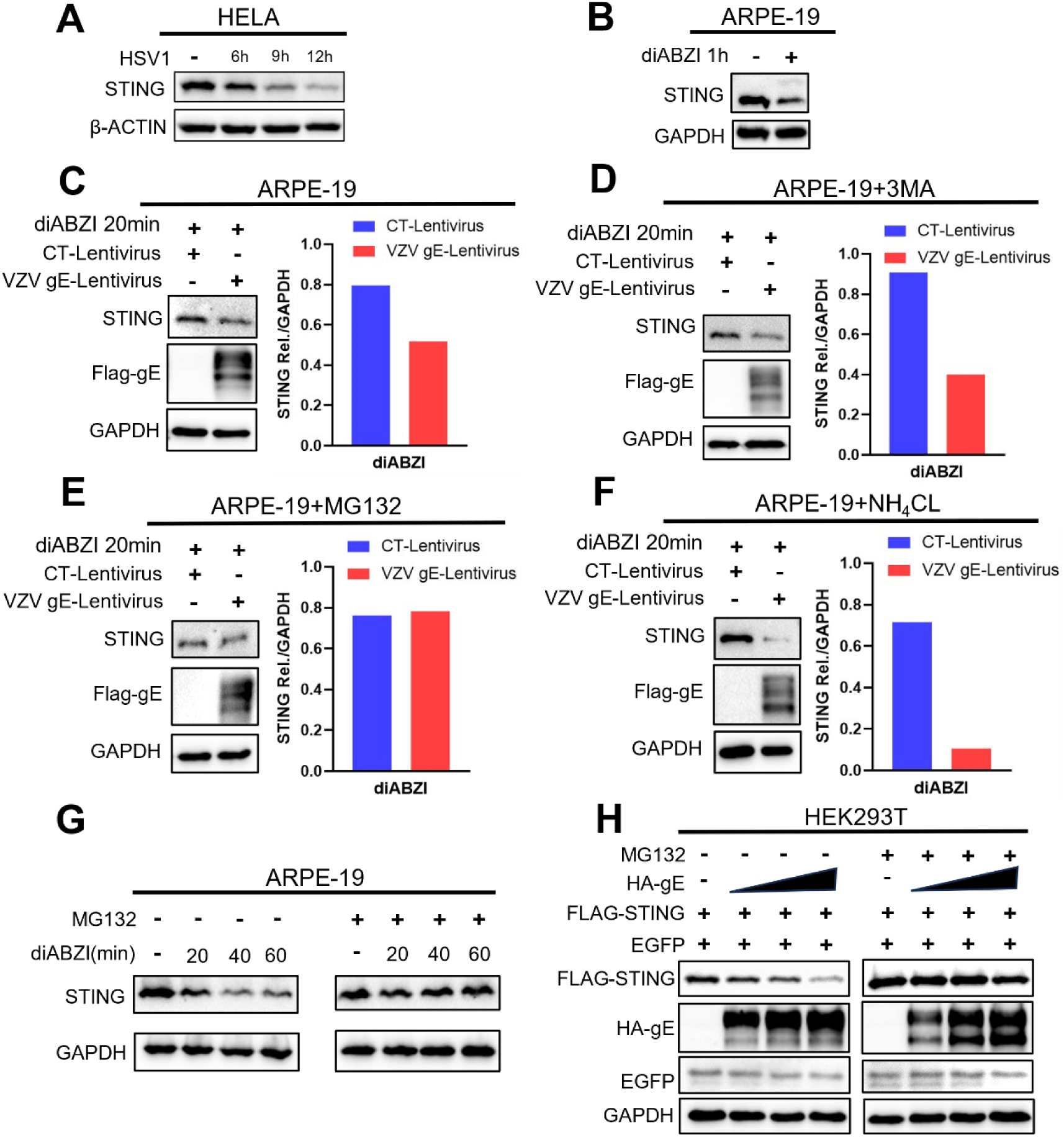
The proteasome inhibitor MG132 can inhibit the degradation of STING that promoted by gE. A: WB results for HELA cells uninfected and infected with HSV1 at 6, 9, and 12 hours. B: WB results for ARPE-19 cells untreated and treated with diABZI (5 μM, 1 hour). C, D, E, F: Sequentially, WB and densitometry results for ARPE-19 cells untreated, treated with 3MA (50 μM, 12 hours), MG132 (5 μM, 12 hours), and NH_4_CL (10 mM, 12 hours) under conditions with diABZI (5 μM) for 20 minutes. G: WB results for ARPE-19 cells untreated and treated with MG132 (5 μM, 12 hours) under conditions with and without diABZI (5 μM) at 20, 40, and 60 minutes. H: The WB data were obtained 24 hours post-transfection in 293T cells with FLAG-STING, EGFP, and a range of concentrations of HA-gE plasmids. Throughout this interval, the experimental conditions encompassed both untreated controls and groups treated with MG132 (at a concentration of 5 μM for 12 hours).

In order to determine the modulatory role of gE on STING’s functionality through its degradation, we screened a panel of inhibitors, included 3MA (a autophagosome inhibitor, Figure 5D), MG132 (a proteasome inhibitor, Figure 5E) and NH4CL (a lysosomal inhibitor, Figure 5F). After STING activation with diABZI treatment, compared with CT-expressing cells, the degradation of STING was increased in gE- expressing cells (Figure 5C). Notably, compared with CT-expressing cells, 3MA (Figure 5D) and NH4CL (Figure 5F) had no affect the degradation level of STING by gE-expressing cells. However, MG132, synchronized STING degradation in both control and gE-expressing cells, suggesting the proteasome’s central role in this process. Our subsequent experiments brought to light that MG132 potently mitigated the degradation of STING in response to diABZI (Figure 5G).

Further experiments indicated that the degradation level of STING was increased along with gE-expressing in a does-dependent manner, but MG132 blocked this effect, implicating the proteasome pathway in gE-induced STING degradation (Figure 5H). We hypothesized that gE suppresses the cGAS-STING pathway by promoting the proteasome pathway to degrade STING.

### gE had no influence in the level of ubiquitination of STING

The proteasome, a colossal protein complex resident within the cell, serves as a central hub for the degradation of intracellular proteins, playing a pivotal role in maintaining cellular homeostasis [21]. It carries out its function by selectively targeting proteins that have been tagged with Ub, a process integral to the regulation of protein turnover [20]. Firstly, we performed a series of gradient transfection with the FLAG- gE plasmid in SHSY5Y cells and examined the ubiquitination level of STING, The results showed that gE had no affect the level of ubiquitination of STING in does-dependent manner (Figure 6A). Similarly, our results also showed that: compared with CT-expressing cells, gE-stable expressing SHSY5Y cells had no affect the level of ubiquitination of endogenous STING even treated them with MG132 (Figure 6B). Therefore, gE may exert its regulatory influence by an Ub-independent proteasome-mediated degradation of STING.

**Figure 6.**
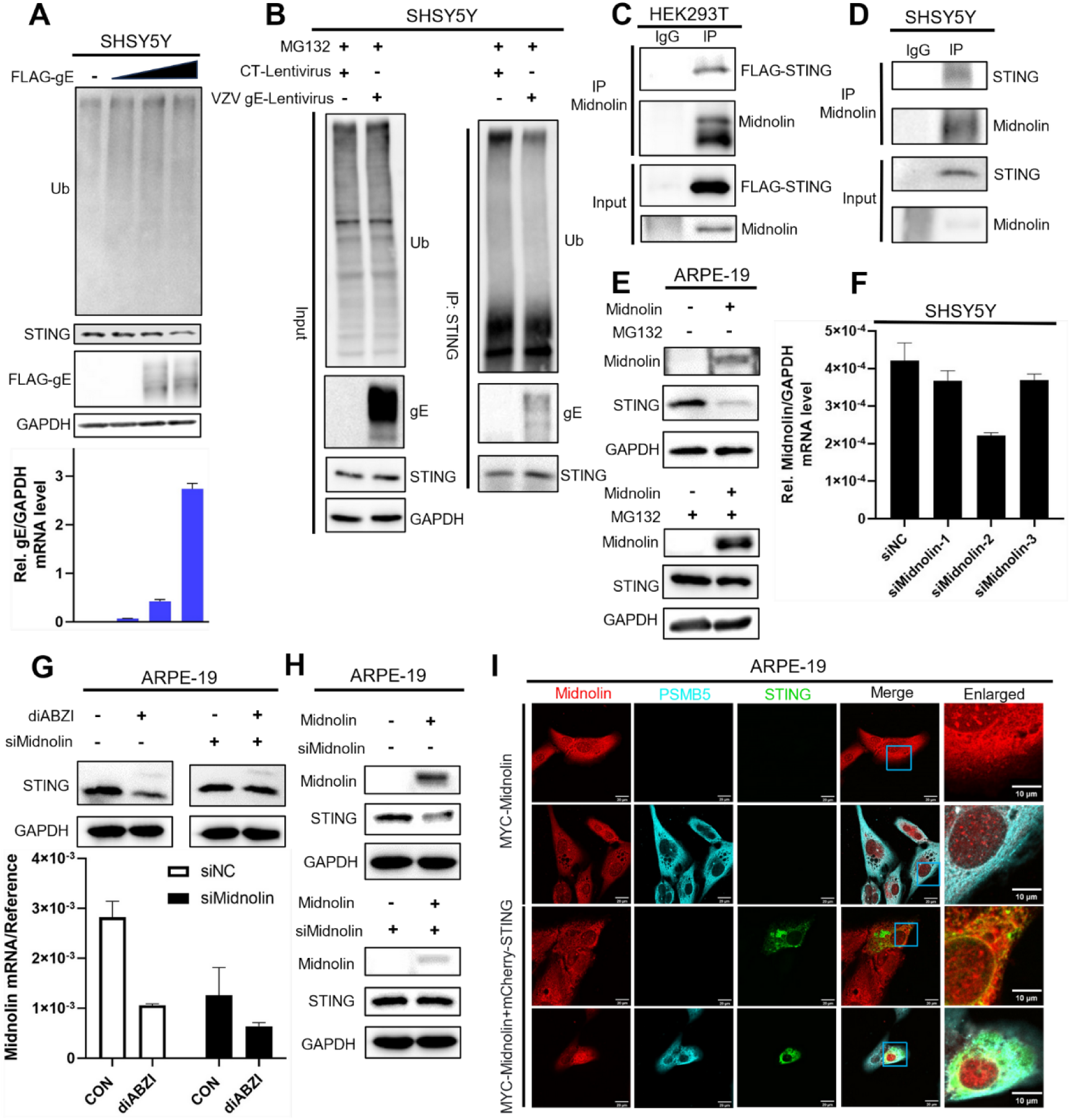
The ubiquitination level of STING is not affected by gE, but STING can be degraded though ubiquitin-independent Midnolin-proteasome pathway. A: The WB data and gE relative mRNA results were obtained 24 hours post-transfection in SHSY5Y cells with a range of concentrations of FLAG-gE plasmids. B: WB results of STING ubiquitination pull-down assays to detect the ubiquitination levels of STING in SHSY5Y-CT and SHSY5Y-gE cells under the treatment with MG132 (5 μM, 12 hours). C: WB results of CO-IP assays for Midnolin and FLAG-STING in HEK293T cells. D: WB results of CO-IP assays for Midnolin and STING in SHSY5Y cells after treatment with MG132 (5 μM, 12 hours). E: WB results for ARPE-19 cells untreated and transfected with Midnolin plasmids for 12 hours, followed by untreated condition and treatment condition with MG132 (5 μM, 12 hours). F: Three Midnolin siRNAs were synthesized, and their interference results were verified in SHSY5Y cells. G: WB and quantitative PCR results for ARPE-19 cells untreated and transfected with siMidnolin-2 for 24 hours, followed by untreated conditions and treatment with diABZI (5 μM, 1 hour). H: WB results for ARPE-19 cells untreated and transfected with Midnolin plasmids for 6 hours, followed by untreated conditions and treatment with siMidnolin-2 for 24 hours. I: MYC-Midnolin and mCherry-STING were co-transfected into 35 mm confocal dishes, MYC-Midnolin (red) and PSMB5 (Proteasome subunit beta type-5, cyan) mCherry-STING (green) were facilitating imaging using confocal microscopy.

### STING can be degraded though ubiquitin-independent Midnolin-proteasome pathway

The Midnolin-proteasome pathway, a novel cellular strategy for protein degradation, eschews the conventional reliance on ubiquitination, providing an expedited route for protein breakdown [22]. Our study explored this pathway’s role in STING degradation. Firstly, we proved that Midnolin could interact with STING both in HEK293T (Figure 6C) and SHSY5Y cells (Figure 6D). Further experiments showed a decrease in STING protein level with Midnolin overexpression in ARPE-19 cells, which can be inhibited by proteasome inhibitor MG132, indicating that Midnolin was involved in STING degradation (Figure 6E). To further investigate Midnolin’s effect on STING, we developed siRNAs to knock-down Midnolin, with siMidnolin-2 being the most effective (Figure 6F). The result showed that the degradation of STING was increased after diABZI treatment, but when we konck-down the expression of endogenous Midnolin, the degradation of STING was restored after diABZI treatment (Figure 6G). Next, we over-expressed Midnolin in ARPE-19 cells, the degradation of STING was increased, but when we konck-down the expression of Midnolin, the degradation of STING was restored (Figure 6H). Confocal microscopy results showed colocalization between MYC-Midnolin, Proteasome subunit beta type-5 (PSMB5) and mCherry-STING, which confirmed Midnolin and STING could co localize in the proteasome (Figure 6I). The above results indicated that Midnolin-proteasome pathway plays a role in STING degradation.

### gE promotes the degradation of STING through the Midnolin-proteasome pathway

To explore how gE influences STING and Midnolin. Firstly, we examined the effect of gE on the expression of Midnolin, the result showed that compared with control cells, the mRNA level of Midnolin was increased under both untreated and daABZI-treated gE-expressing cells (Figure 7A). Next, CO-IP assays confirmed that endogenous Midnolin could interact with gE both in HEK293T cells (Figure 7B) or SHSY5Y stable expressed gE cells (Figure 7C). Further result showed that gE promoted the interaction between STING and Midnolin (Figure 7D). To further investigate the effect of glycosylation of gE on the interaction between STING and Midnolin, we treated cells with tunicamycin, the results showed that deglycosylated gE significantly promoted the interaction between STING and Midnolin (Figure 7E). Experiments on SHSY5Y-gE cells revealed a marked enrichment of Midnolin protein within the STING in cells harboring gE, a phenomenon that become more apparent in the treatment of diABZI-induced pathway activation (Figure 7F). Intriguingly in SHSY5Y-gE cells, the introduction of siMidnolin reversed the gE-promoted degradation of STING, regardless of whether the cGAS-STING signaling pathway was activated by diABZI (Figure 7G). This suggests that the role of gE extends beyond merely transcriptionally upregulating Midnolin mRNA; it also enhances the process of STING degradation via Midnolin.

**Figure 7.**
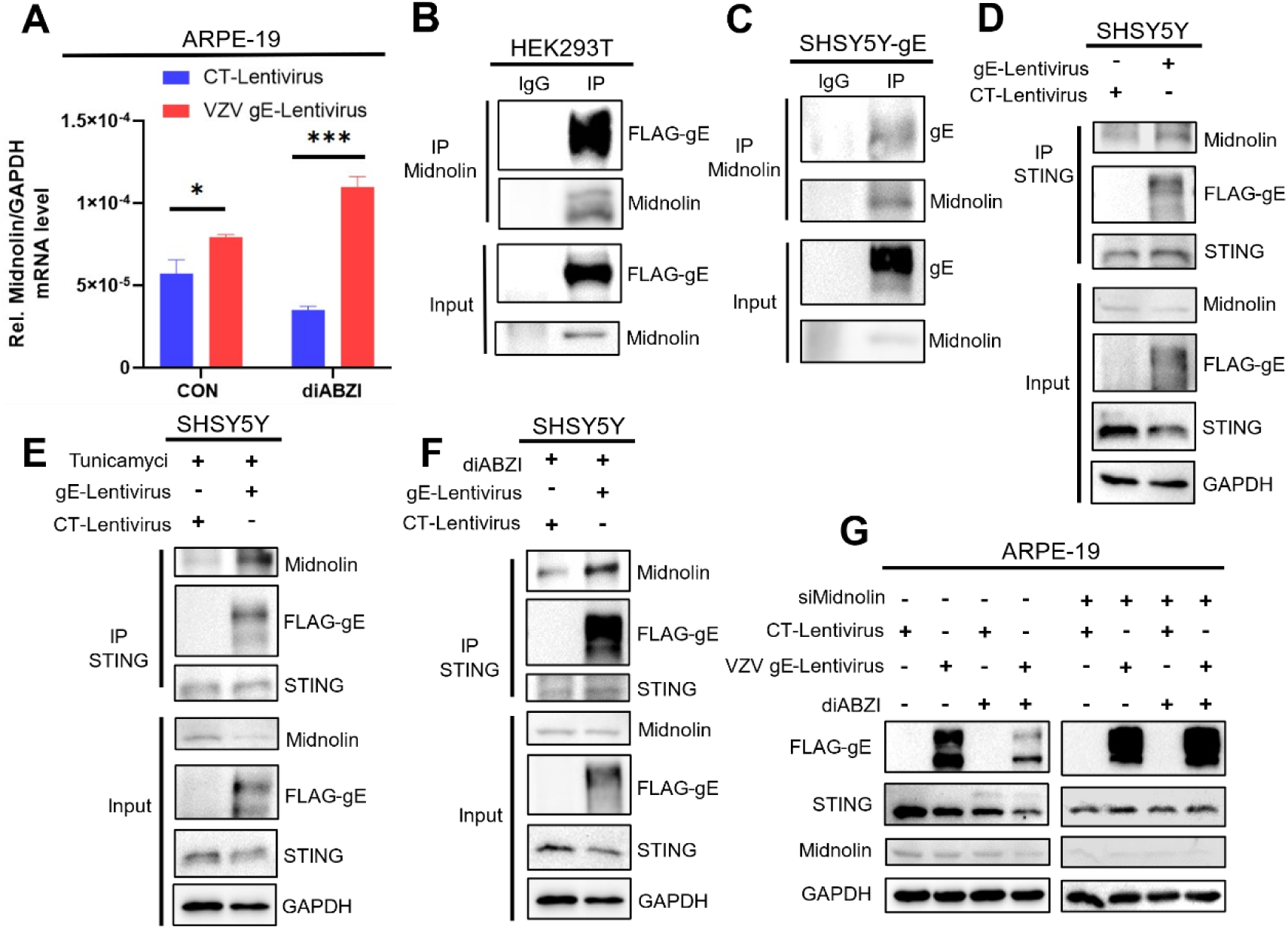
gE promotes the degradation of STING through the Midnolin-proteasome pathway. A: Results of Midnolin relative mRNA levels in ARPE-19-CT and ARPE-19-gE cells after untreated conditions and treatment with diABZI (5 μM, 1 hour). B: WB results of CO-IP assays for Midnolin and FLAG-gE in HEK293T cells. C: WB results of CO-IP assays for Midnolin and gE in SHSY5Y-gE cells following treatment with MG132 (5 μM, 12 hours). D: WB results of CO-IP assays for Midnolin and STING in SHSY5Y-CT and SHSY5Y-gE cells. E: WB results of CO-IP assays for Midnolin and STING in SHSY5Y-CT and SHSY5Y-gE cells with treatment of tunicamycin (2 μg/mL for 12 hours). F: WB results of CO-IP assays for Midnolin and STING in SHSY5Y-CT and SHSY5Y-gE cells under treatment of diABZI (5 μM, 1 hour). G: WB results in ARPE-19-CT and ARPE-19-gE cells without or after transfection with siMidnolin-2 for 24 hours under untreated conditions and treatment of diABZI (5 μM, 1 hour).

### gE inhibits cGAS-STING-triggered signaling pathway in mouse

To study gE’s role in the cGAS-STING pathway in vivo, we injected C57BL/6 mice with AAV-CT or AAV-gE via tail vein and infected them with HSV1 after 21 days (Figure 8A). qRT-PCR analysis of blood samples showed that compared to control group, the mRNA levels of IFN-β, ISG15 and IFIT1 was increased in HSV1 infected group, compared to AAV-CT group infected with HSV1, the mRNA levels of IFN-β, ISG15 and IFIT1 was decreased in AAV-gE group infected with HSV1 (Figure 8B), suggesting gE dampens the immune response. However, when we pre-treated with the STING inhibitor H151, compared to AAV-CT group, the mRNA levels of IFN-β, ISG15 and IFIT1 was no influence in AAV-gE group infected, suggested that gE inhibited the immune response through regulating STING triggered signaling pathway in vivo (Figure 8B). HE staining of liver tissue revealed typical hepatocyte arrangement in the Mock group, while the HSV1 group showed significant inflammatory cell infiltration or tissue edema. The AAV-gE group had a more intense inflammatory response than AAV-CT post-HSV1 infection, but H151 treatment mitigated this difference between AAV-CT group and AAV-gE group (Figure 8C). Western blot further confirmed that AAV-gE group reduced HSV1-induced pIRF3 activation compared to AAV-CT group, but H151 abolished this effect between AAV-CT group and AAV-gE group (Figure 8D).

**Figure 8.**
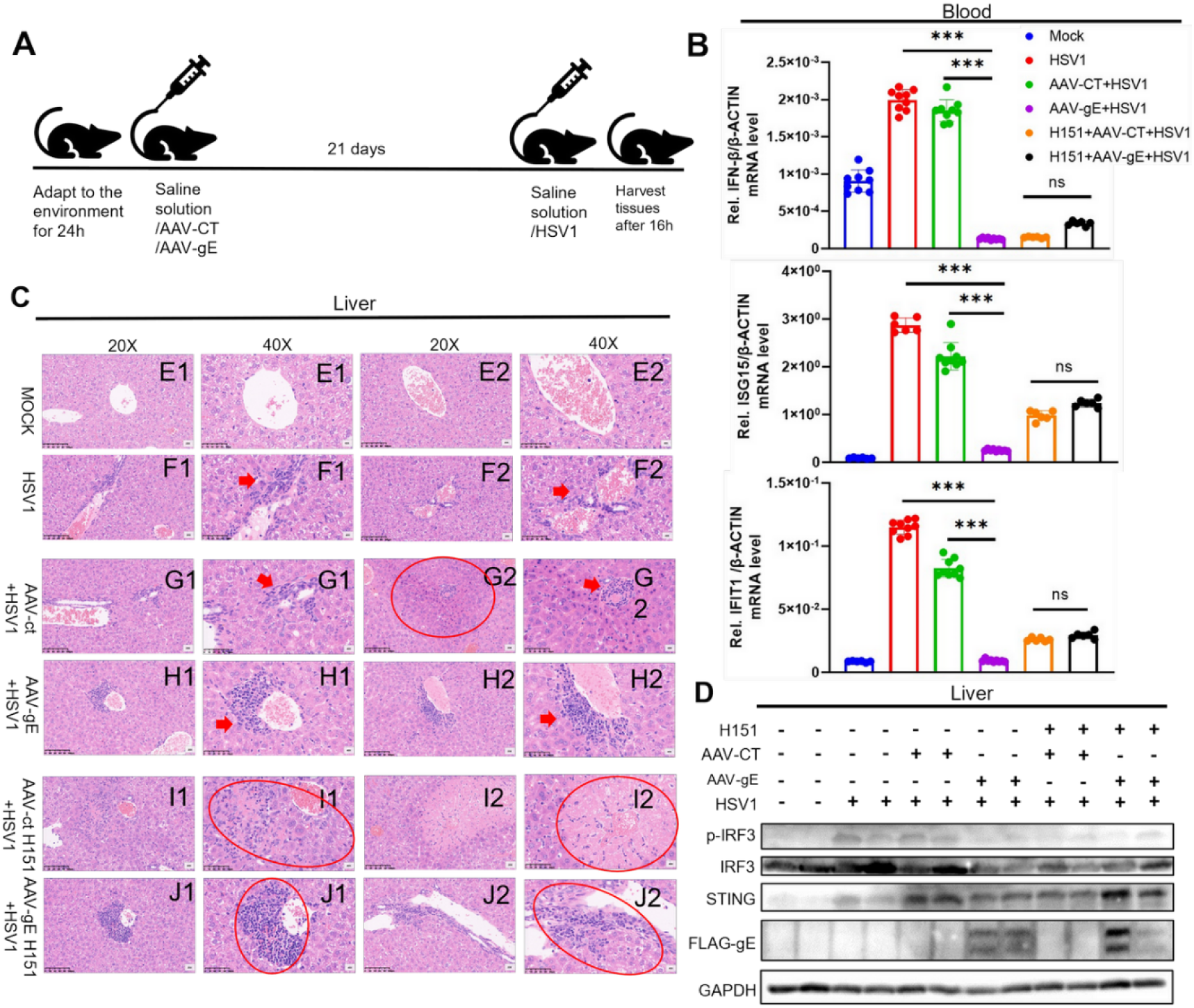
gE inhibits cGAS-STING-triggered signaling pathway in mouse. A: A flowchart of the animal experiment is presented, where C57BL/6 mice were divided into 6 groups (8-9 mice per group). The specific groups were as follows: Mock, HSV1, AAV-CT+HSV1, AAV-gE+HSV1, AAV-CT+H151+HSV1, and AAV-gE+H151+HSV1. H151 was administered via intraperitoneal injection at a dosage of 25 mM per mouse, which is equivalent to 0.32 mg/kg. B: The relative mRNA levels of IFN-β, IFIT1, and ISG15 in the ocular blood of mice from each group are shown. C and D: Hematoxylin and Eosin staining and WB results of the livers from mice in each group are depicted.

The above results demonstrated that the incompletely glycosylated VZV gE can bind to STING, which may promote the degradation of STING through the ubiquitin-independent Midnolin proteasome pathway and downregulate the release of anti-viral IFNs and cytokines, thereby facilitating VZV proliferation (Figure 9).

**Figure 9.**
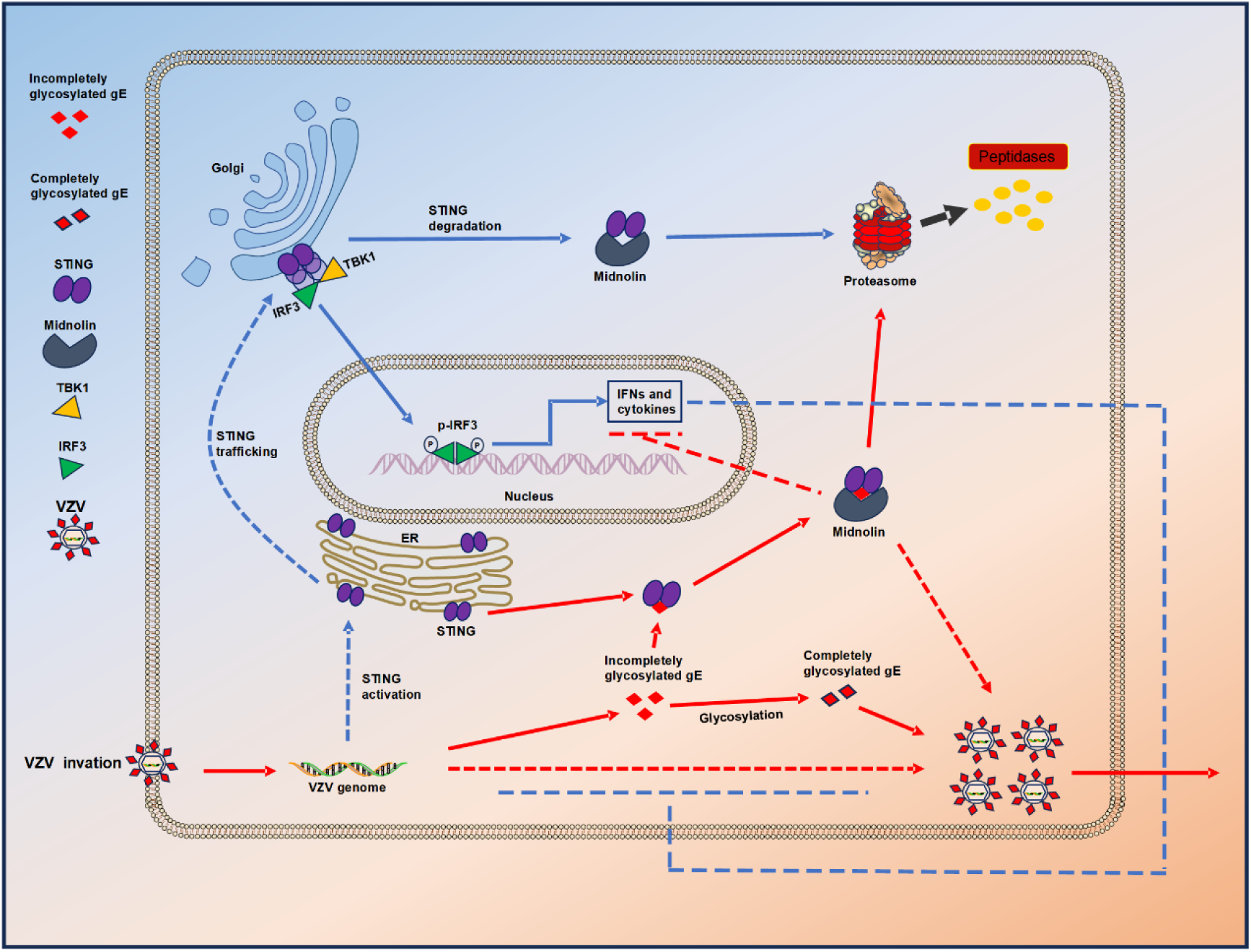
A proposed model for the regulation of gE inhibits cGAS-STING-triggered signaling through Midnolin-proteasome pathway. When the cGAS-STING pathway is activated by stimulation, anti-viral IFNs and cytokines are released. However, in VZV infection, the incompletely glycosylated VZV gE can bind to STING, which may promote the degradation of STING through the ubiquitin-independent Midnolin proteasome pathway and downregulate the release of anti-viral IFNs and cytokines, thereby facilitating VZV proliferation.

## Discussion

The cGAS-STING signaling pathway is pivotal in innate immunity, serving as a critical pattern recognition and effector mechanism. However, hyperactivation triggered by the detection of pathogen or aberrant self-DNA can lead to excessive Type I IFN production, potentially precipitating a range of inflammatory diseases, including inflammatory bowel disease, cardiovascular disorders, and cancers characterized by chronic inflammation [15] [28] [29] [30] [31] [32]. Consequently, prompt STING degradation is crucial not only to mitigate exaggerated immune responses but also to maintain intracellular homeostasis, thereby preventing autoimmune disorders. Our groundbreaking research unveils a novel mechanism by which Midnolin facilitates the post-activation degradation of STING, a process that occurs through a newly identified ubiquitin-independent pathway mediated by the Midnolin-proteasome complex [22]. Furthermore, we have discovered that the under-glycosylated form of gE amplifies STING degradation via the Midnolin pathway, resulting in the suppression of IFN-β- related gene expression. This, in turn, enables immune evasion and augments the proliferation of VZV.

The dynamic regulation of STING’s activity is essential for maintaining intracellular homeostasis, as its activation simultaneously triggers a sequence leading to its deactivation [33]. Various mechanisms have been identified to facilitate the prompt degradation of STING upon activation, including trafficking to Rab7-positive endolysosomes [19], ESCRT-dependent redirection to endolysosomal compartments [34], autophagy [35] [36], and the ubiquitin-proteasome system [37]. When STING is inactive, it undergoes ER-associated degradation (ERAD) [38], limiting its activation potential and preserving cellular homeostasis. This delicate balance underscores the sophistication of cellular mechanisms governing the lifecycle of critical immune regulators like STING. Midnolin has been reported to rapidly degrade proteins belonging to the IEG family, thereby enabling the swift control of the signal transduction network regulated by IEG expression [22]. This ensures that following transient activation, the network can be promptly modulated to prevent the occurrence of diseases such as cancer and immune deficiencies, which may arise from dysregulation [22]. Similarly, the cascade-amplified antiviral response triggered by STING activation also necessitates such rapid control. The Midnolin-mediated rapid degradation of STING may offer novel research perspectives for diseases associated with excessive STING activation.

A recent study conducted by Oh SJ and his team [39] uncovered that gE interacts with LC3, thereby facilitating mitophagy via the PINK1/Parkin pathway and impeding the trafficking of STING from the endoplasmic reticulum to the Golgi apparatus. Extending these findings, our research delves deeper into the interaction between the under-glycosylated variant of gE and STING. It is remarkable that prior research has established an interaction between non-glycosylated gE and IDE within the endoplasmic reticulum [11], suggesting a potential ERAD mechanism for IDE that aligns with our observations. Specifically, we have noted that the complex formed by under-glycosylated gE and STING may trigger the ERAD degradation pathway due to its misfolded structure. However, a pressing question for further exploration is whether Midnolin also plays a role in the ERAD process. Additional investigation is necessary to solidify this connection.

The extracellular N-terminal domain of gE derived from VZV comprises a substantial sequence of approximately 500 amino acids, posing a formidable obstacle for structural elucidation due to its considerable size and intricate nature. This domain plays a pivotal role in VZV proliferation, as mutations within the amino acid segment ranging from 27 to 90 disrupt gE’s interaction with IDE [7], while the cysteine-rich region spanning amino acids 208 to 236 is indispensable for forming a natural heterodimer with gI [40]. Our research further unveiled that the amino acid segment from 350 to 480 of gE interacts with STING, resulting in incomplete glycosylation of gE. A notable limitation here is that commercial antibodies are unable to detect the incompletely glycosylated form of gE [11]. Therefore, to gain a deeper understanding of the interaction between gE and STING, it may be necessary to adopt further research strategies, such as modifying the glycosylation sites within the full-length gE protein of the VZV virus or developing antibodies that can recognize a variety of glycosylation patterns of gE.

Moreover, the amino acid sequences located within the 350 to 480 range in the gE- STING interaction domain have emerged as promising candidates for the development of potent antiviral therapies. By precisely engineering peptides targeted to this specific domain, we may potentially hinder the proliferation of the VZV virus. Additionally, a recent study [14] has underscored the remarkable antigenic properties of a peptide encompassing amino acids 31 to 358 within gE, highlighting its substantial potential for the creation of a vaccine against shingles. However, given the overlap between this peptide and the STING interaction domain, and the presence of an incompletely glycosylated domain within gE, further research is urgently required to strike a harmonious balance between antiviral therapies and vaccine development, which is crucial for the effective management of viral infections.

## Materials and Methods

### Ethics statements

All animal experiments were conducted in strict adherence to the ethical guidelines outlined in the Animal Welfare Act and the National Institutes of Health’s directives for the treatment of laboratory animals in scientific research. The protocols involving mice were meticulously reviewed and granted approval by the Institutional Animal Care and Use Committee of the Laboratory Animal Science Institute at Jinan University, with the assigned ethical committee approval number IACUC-20210429-01.

### Animals and animal experimental procedure

Four-week-old male C57BL/6 mice, weighing 17 to 18 grams, were obtained from the Guangdong Provincial Medical Laboratory Animal Center. We used a specially tailored rAAV2/9 serotype virus from Heyuan Biotech, incorporating the gE gene sequence from VZV, known for its tropism towards various tissues. The virus was administered at a high titer, with each mouse receiving 1x10^11^ V.G. via tail vein injection. The mice were divided into six groups for the experiment: a mock group, an HSV1-only group, and four groups receiving different combinations of AAV-CT or AAV-gE, with or without HSV1, and H151. After a 21-day observation period, mice were injected with HSV1 (2x10^7^ PFU per mouse) and, for the H151 treatment group, the inhibitor was given 2 hours prior at a dosage of 0.32 mg/kg. At 16 hours post-infection, mice were euthanized for further analysis, including Western blot, qPCR, and HE staining.

### Cell lines and cultures

The human embryonic kidney cell line (HEK293T), African green monkey kidney epithelial cell line (Vero), U-251MG cell line (U251) and the human cervical carcinoma cell line (HELA) were procured from the American Type Culture Collection. The human Neuroblastoma SH-SY5Y cell line (SHSY5Y) and the Adult Retinal Pigment Epithelial 19 cell line (ARPE-19) were graciously provided by Professor Zhu Hua’s research group [41]. Cells were cultivated in Dulbecco’s Modified Eagle Medium (DMEM). Each medium was enriched with 10% fetal calf serum, 100 U/mL penicillin, and 100 μg/mL streptomycin, and the cells were kept at 37 °C in an incubator with a 5% CO2 atmosphere. Following this, the cells were transferred to fresh medium and incubated for 24 hours before undergoing various treatments. Subsequently, the cells were harvested for qRT-PCR and immunoblot analysis.

### Viruses and Plaque assay

The HSV-1 strain was a gift from Dr. Bo Zhong of Wuhan University. Vero cells were grown in DMEM with 10% fetal calf serum and antibiotics at 37°C. Viral stocks were produced by infecting Vero cells at an MOI of 0.03 for 36 hours, and titers were determined by plaque assay in Vero cells. Control mock-infected cells were treated similarly without virus.

The VZV-GFP strain, a derivative of the rOka vaccine strain with GFP and luciferase genes, was provided by Zhu Hua’s research group, which exhibits growth characteristics comparable to the original Oka strain [26]. The viral titer was determined using an infectious focus assay. ARPE-19 cells were infected with diluted VZV-GFP suspension, and plaques were counted under fluorescence microscopy at 72 hours post-inoculation. Given the highly cell-associated nature of VZV under tissue culture conditions, cells with virus were stored in liquid nitrogen after viral titers were assessed. Luciferase activity was measured using a Luciferase Assay System (Promega, E4030).

### Plasmids Constructs and transfection

Our laboratory previously constructed plasmids including pcDNA3.1(+)- 3×FLAG-Vector/IRF3/TBK1/cGAS and pcAGGS -HA-Vector/STING. Utilizing the ClonExpress MultiS One Step Cloning Kit (Vazyme, C113-01), we cloned the 10 glycoprotein genes of VZV— encompassing ORFS/L (ORF0), gK (ORF5), gN (ORF9A), gC (ORF14), gB (ORF31), gH (ORF37), gM (ORF50), gL (ORF60), gI (ORF67), and gE (ORF68) — into the pcDNA3.1(+) expression vector. The PCR products for the truncated forms of gE were inserted into various restriction enzyme sites of the pcDNA3.1(+)–3× FLAG plasmid. The EGFP-Vector and pCMV-mCherry-Vector were kindly provided by Professor Chen Tongsheng’s research group from the School of Life Sciences, South China Normal University. Primer sequences are detailed in Supplementary Table 1.

### Coimmunoprecipitation (CO-IP), immunoblot assays and western blot (WB) analysis

#### CO-IP and immunoblot assays

Whole-cell lysates from HEK293T or SHSY5Y cells were prepared using a lysis buffer containing 50 mM Tris-HCl at pH 7.5, 300 mM NaCl, 1% Triton-X, 5 mM EDTA, and 10% glycerol. The cells were lysed, and the lysates were subjected to immunoprecipitation using immunoglobulin G (IgG) from Invitrogen, anti-FLAG, anti-HA antibodies, or specific target antibodies. After rotating the mixture at 4 °C for 30 minutes, the lysates were centrifuged at 13,500 × g for 10 minutes to pellet any cellular debris. A portion of the supernatant was reserved as the input control, while the remainder was incubated with the respective antibodies overnight at 4 °C. This was followed by the addition of Protein G Sepharose beads from GE Healthcare and a further 2-hour incubation at 4 °C. The immunoprecipitates were then thoroughly washed four to six times with the same lysis buffer used initially, boiled in protein loading buffer, and subsequently analyzed by WB to assess the protein interactions.

#### Western blot analysis

Cells or animal tissuewere harvested and lysed in a buffer solution containing 50 mM Tris-HCl at pH 7.4, 300 mM NaCl, 1% Triton X-100, 5 mM EDTA, and 10% glycerol. Prior to lysis, a protease inhibitor cocktail (10%, Roche, catalog number 04693116001) was incorporated into the buffer to prevent protein degradation. The concentration of proteins in the lysates was determined using the Bradford assay, as per the manufacturer’s instructions (Bio-Rad, Richmond, CA). Equal amounts of protein (50 μg) from each cell lysate were loaded and separated by SDS-polyacrylamide gel electrophoresis (PAGE) on gels with concentrations ranging from 8 to 12%. The resolved proteins were then transferred onto nitrocellulose membranes (Amersham, Piscataway, NJ). To block nonspecific binding sites, the membranes were immersed in 5% skim milk for 2 hours. Following the blocking step, the nitrocellulose membranes were rinsed three times with PBS containing 0.1% Tween 20 and then incubated with the appropriate primary antibodies. After incubation with the primary antibodies, the membranes were washed again and incubated with the corresponding secondary antibodies. The protein bands were finally visualized using a Bio-Rad Image Analyzer (733BR3722).

#### Antibodies and reagents

Dulbecco’s Modified Eagle Medium (DMEM) was sourced from Gibco. Reagents such as 3-MA (M9281) and NH_4_CL (326372) were acquired from Sigma. Additional compounds, including MG132 (S2619) and diABZI (S8796), were procured from Selleck. Tunicamycin (HY-A0098) was obtained through Medchemexpress, and the Lipofectamine 2000 transfection reagent (11668019) was secured from InvivoGene. Antibodys against FLAG (F3165), HA (H6908),MYC (C3956) and monoclonal mouse anti-GAPDH (G9295) were purchased from Sigma. Monoclonal rabbit anti-β-Actin (AC026) were purchased from ABclonal. Ubiquitin mouse mAb (P4D1), monoclonal rabbit anti-STING (D2P2F), monoclonal Anti-IRF3 (D6I4C), monoclonal Anti-Phospho-IRF3(Ser396)(4D4G), monoclonal Anti-TBK1(D1B4), monoclonal Anti-Phospho-TBK1 (Ser172) (D52C2) were purchased from Cell Signaling Technology. Monoclonal Anti-VZV gE (ab272686) was purchased from Abcam. Polyclonal antibody anti-Midnolin (18939-1-AP) and polyclonal antibody anti-PSMB5 (19178-1-AP) were purchased from Proteintech.

### Generation of stable cell lines using lentivirus or retrovirus

HEK293T cells were plated in 10 cm culture dishes and subjected to transfection with either the plenti-3×FLAG-gE construct or a control empty vector, in conjunction with the packaging plasmids psPAX2 and pMD2G, utilizing Lipo2000 as the transfection reagent. The culture medium was refreshed 12 hours post-transfection. At 36 and 60 hours following transfection, the cell supernatants, which contained the lentivirus, were harvested and passed through a 0.45 μm filter to remove any particulates. Subsequently, SHSY5Y, ARPE-19, U251, and HELA cells were infected with the collected lentivirus for a duration of 24 hours in the presence of polybrene (Sigma, TR-1003) to enhance the infection efficiency. Forty-eight hours after the initiation of the infection process, the cells were subjected to selection with puromycin (Sigma, P8833) for a period of 4 to 7 days to isolate successfully transduced cells. A subset of these cells was then harvested for further characterization by WB analysis to confirm the expression of the introduced construct.

To generate a stable STING-knockout (KO) cell line, the Lenti CRISPR v2 puro-sgRNA was employed, alongside the packaging plasmids psPAX2 and pMD2G. The cell supernatants containing the lentiviral particles were collected and used to infect SHSY5Y cells following the same protocol as for the lentivirus infection. For the design of the sgRNAs targeting the STING gene, two pairs of oligos incorporating the BsmBI restriction enzyme site were utilized, with the sequences as follows: STING sgRNA-1- F: CACCGGGCCGACCGCATTTGGG; STING sgRNA-1-R: AAACCCCAAATGCGGTCGGCCCGC; STING sgRNA-2-F: CACCGTACGCAAGAGTTGCCCGGC; STING sgRNA-2-R: AAACGCCGGGCAACTCTTGCGTAC. These sgRNAs were identified using the guide design resources available on Dr. Feng Zhang’s laboratory website (https://zlab.squarespace.com/guide-design-resources), which offers tools for CRISPR/Cas9 design.

### RNA extract and Real-time PCR

Total cellular RNA was extracted using the Trizol reagent from Invitrogen, following the manufacturer’s protocol. Subsequently, 1 μg of RNA was converted to cDNA in a reaction containing 0.5 μL of oligo (dT) and 0.5 μL of random primers. This reaction was carried out at 37°C for 60 minutes, followed by a denaturation step at 72°C for 10 minutes. The synthesized cDNA served as the template for real-time PCR analysis. The real-time PCR was conducted on a LightCycler 480 thermal cycler from Roche, employing the following thermal profile: initial activation of the polymerase at 95°C for 5 minutes, followed by 45 cycles of denaturation at 95°C for 15 seconds, annealing at 58°C for 15 seconds, and extension at 72°C for 30 seconds. Fluorescence data were collected and analyzed at the extension step. To verify the specificity of the amplification, a final melting curve analysis was performed, ranging from 50°C to 95°C. The primers utilized for the real-time PCR detection are as follows: hGAPDH-f: AAGGCTGTGGGCAAGG, hGAPDH -r: TGGAGGAGTGGGTGTCG; hIFN-β-f: AGTAGGCGACACTGTTCGTG, hIFN-β-r: GCCTCCCATTCAATTGCCAC; hTNF-α-f: CTCTTCTGCCTGCTGCACTTTG, hTNF-α-r: ATGGGCTACAGGCTTGTCACTC; hIL-6-f: AGACAGCCACTCACCTCTTCAG, hIL-6-r : TTCTGCCAGTGCCTCTTTGCTG. hMidnolin-f: GCCTGGCTCTTCTCCACAAA, hMidnolin-r : GAAGTCACTGACCTGCGTCT. VZV-gE-f: GTATCGATAGCGGGGAACGG, VZV-gE-r : TGGCCTTGGGGTTTTGGATT.

### siRNA

Small interfering RNAs (siRNAs) designed to target Midnolin were custom synthesized and introduced into the cells using Lipofectamine 2000 reagent. Post-transfection, the cells were subjected to either quantitative PCR (qPCR) or immunoblot analysis to evaluate the efficacy of siRNA-mediated gene knockdown. The siRNA sequences utilized in this study are listed below: siMidnolin RNA-1: GATGCAAGCTCTCGAGAGT, siMidnolin RNA-2: GCGACCACATGATGTTCGT, siMidnolin RNA-3: TCCTGCAGATCCTGAACGA.

### IFN-β and ISRE reporter assay

HEK293 cells were transfected with a firefly luciferase reporter construct (100 ng) and a TK-driven Renilla luciferase control plasmid (10 ng), along with the plasmids of interest or an empty control vector (100 ng), using a standard calcium phosphate precipitation method. Twenty-four hours post-transfection, luciferase activities were measured using a dual-specific luciferase assay kit from Promega. The firefly luciferase activity was normalized to the Renilla luciferase activity to calculate the relative luminescence units.

### Confocal microscopy

HEK293T were transfected with the respective plasmids for a period of 24 to 36 hours. After washed with PBS, the cells were then examined using a confocal laser scanning microscope (Leica, TCS, SP8) . ARPE-19 were transfected with the respective plasmids for a period of 24 hours. Subsequently, the cells were fixed using an iced methanol at 4 °C for 15 minutes. Following fixation, the cells were rinsed with PBS three times and and finally blocked with PBS containing 5% BSA for 1 h. The cells were then incubated with the Polyclonal antibody anti-Midnolin (18939-1-AP) and polyclonal antibody anti-PSMB5 (19178-1- AP) overnight at 4°C, followed by incubation with FITC-conjugate donkey anti-mouse IgG (Abbkine) and Dylight 649-conjugate donkey anti-rabbit IgG (Abbkine) for 1 h. After washed three more times with PBS, the cells were then examined using the confocal laser scanning microscope.

### Statistics and reproducibility

Each experimental procedure was conducted a minimum of three times, yielding consistent outcomes. For statistical evaluation, the t-test was employed to analyze the differences between two groups, while one-way analysis of variance (ANOVA) was utilized for comparisons among multiple groups, with the aid of SPSS 27.0 software. The results were deemed statistically significant at the following P-value thresholds: * for P ≤ 0.05, ** for P ≤ 0.01, and *** for P ≤ 0.001. The designation ’ns’ indicates no statistical significance.

## Acknowledgments

This work was supported by the National Natural Science Foundation of China (Grant 62275103 to Xiaoping Wang and Grant 32200117 to Pan Pan). Additional funding was provided by the Science and Technology Projects in Guangzhou (Grant 2023A03J1020 to Xiaoping Wang), as well as the Clinical Frontier Technology Program of the First Affiliated Hospital of Jinan University in China (Grant No. JNU1AF-CFTP-2022-a01212 to Xiaoping Wang). Furthermore, the Guangdong Basic and Applied Basic Research Foundation (Grant 2024A1515013063 to Pan Pan).

## Author Contributions

Y.K.L, P.P., and X.P.W. designed research; L.M.C., L.A.D., S.H.N., Y.Z., F.Y.L, X.J.D., C.L.L., M.R.L., Y.Q.L., W.W.G., X.F.L. and Q.Y. performed research; L.M.C. and Z.W.L. analyzed data; X.P.W. revised the paper; L.M.C., P.P., and Y.K.L wrote the paper.

---

**Supplementary Table 1:**
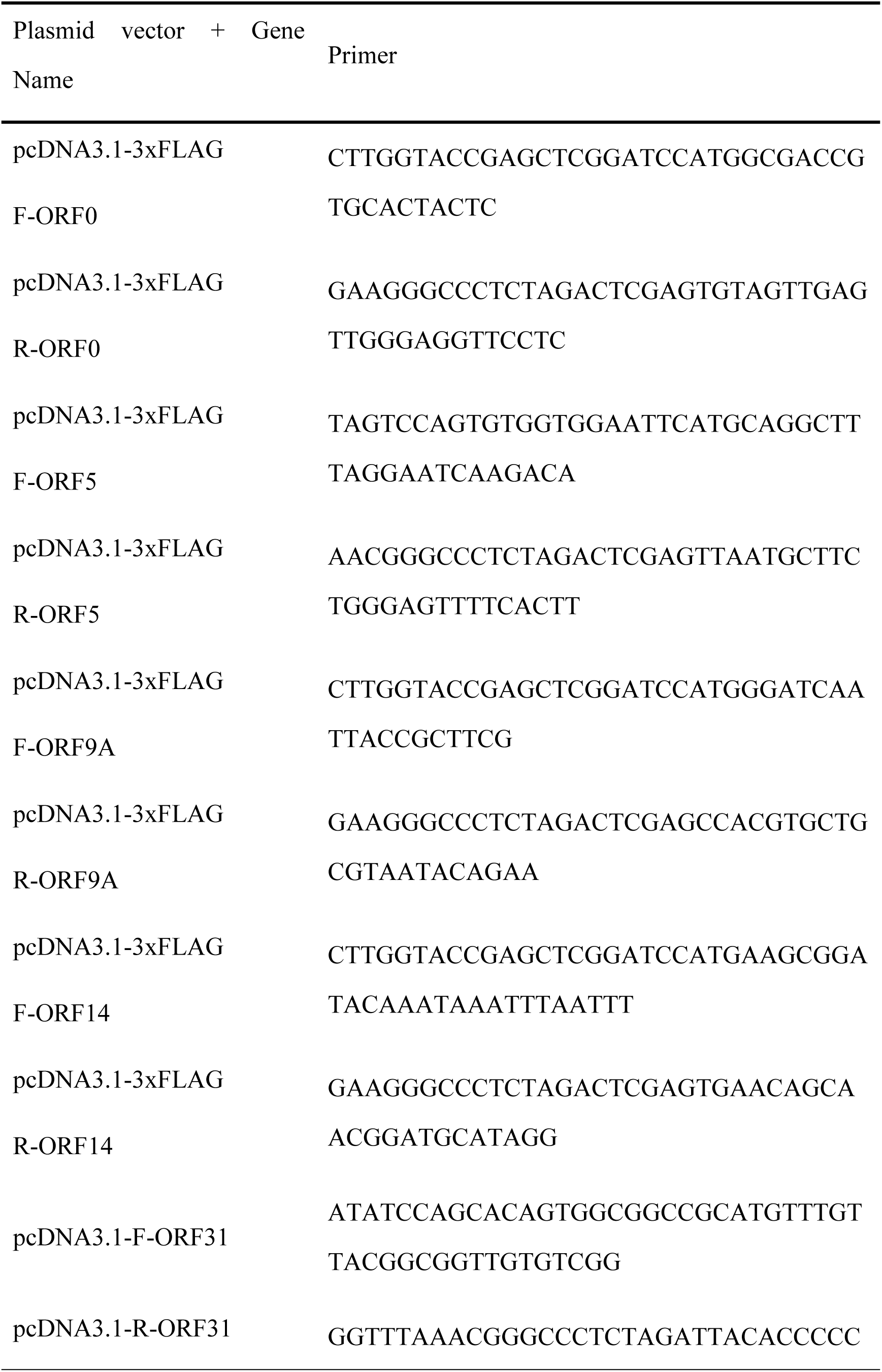
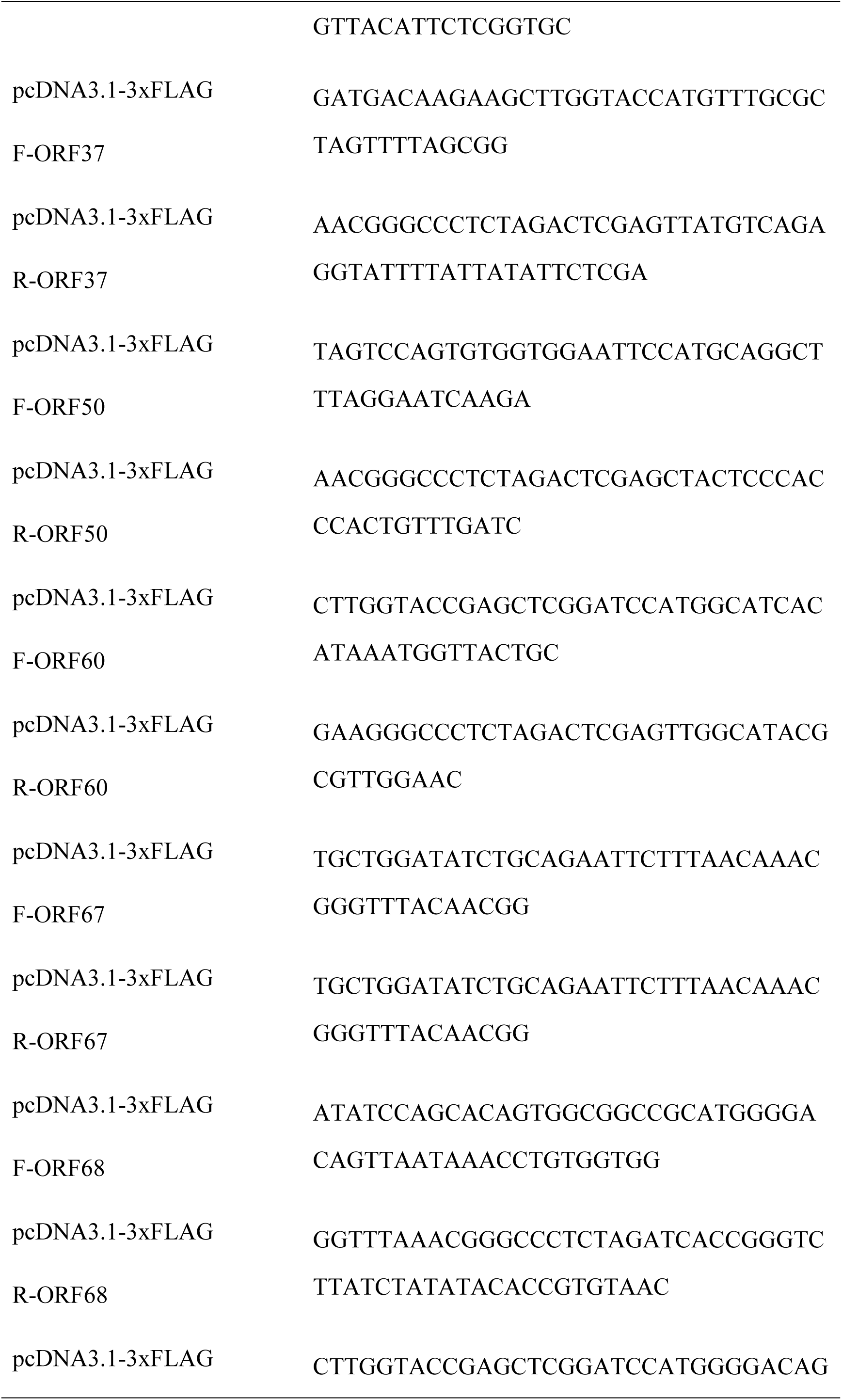
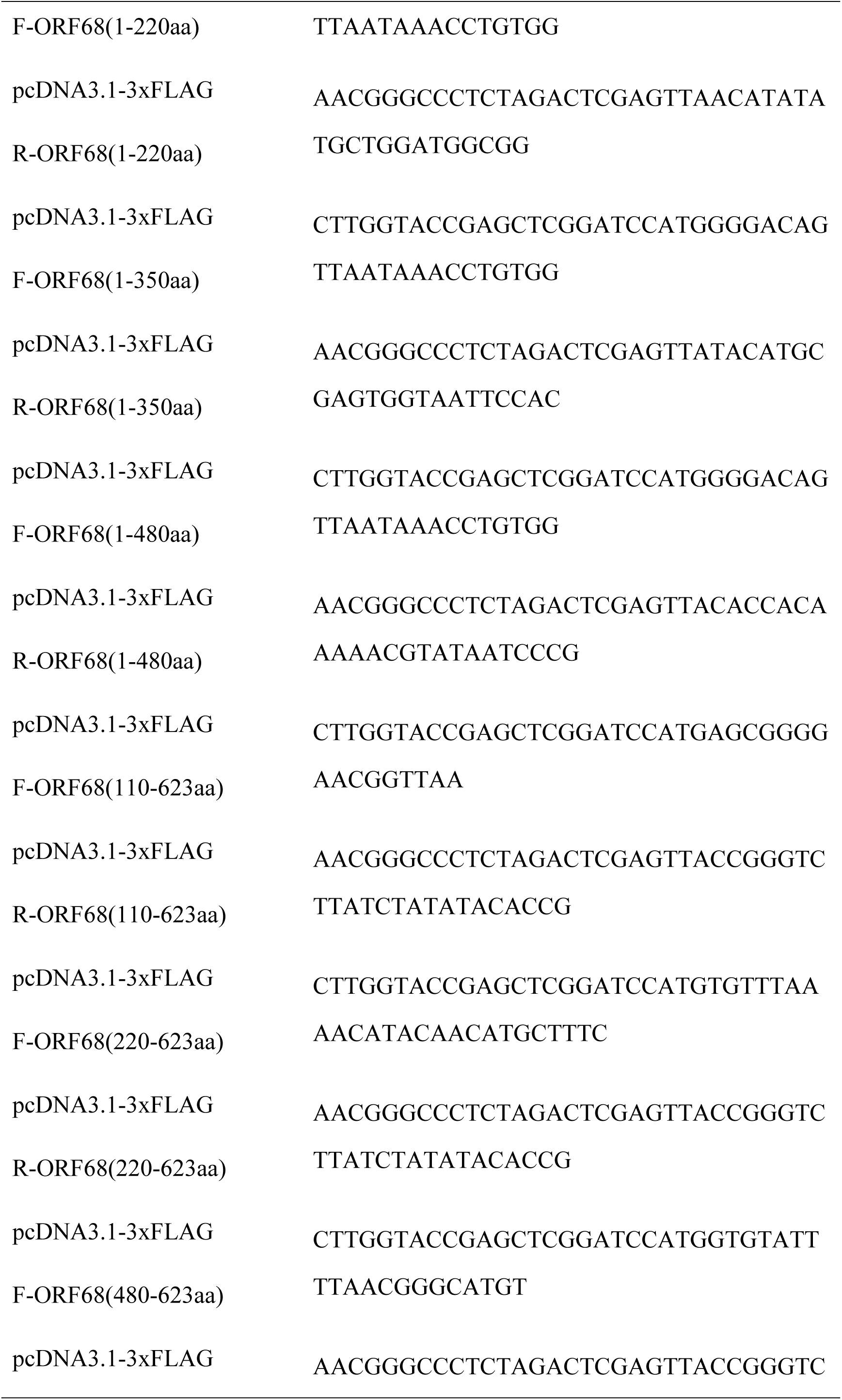
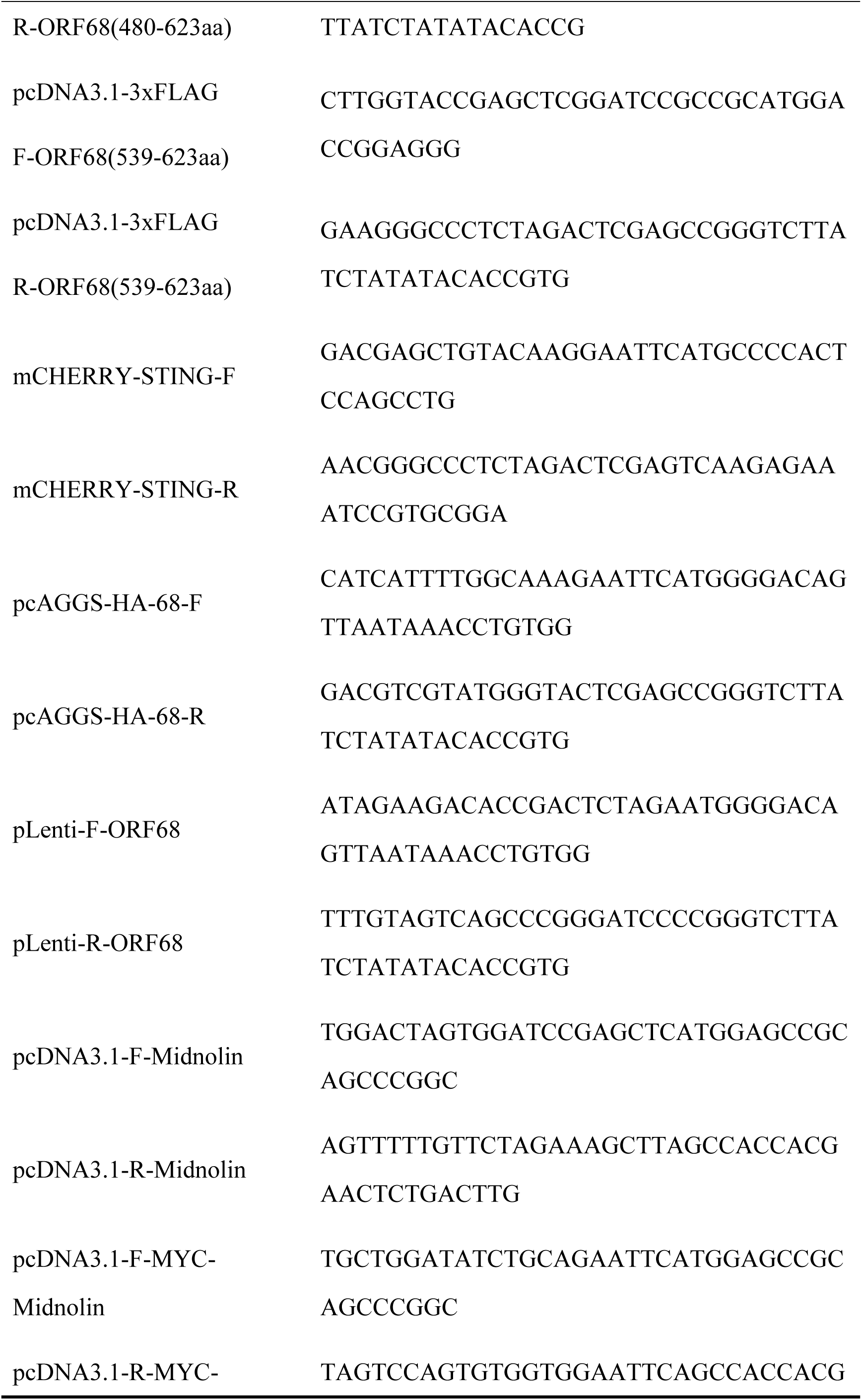
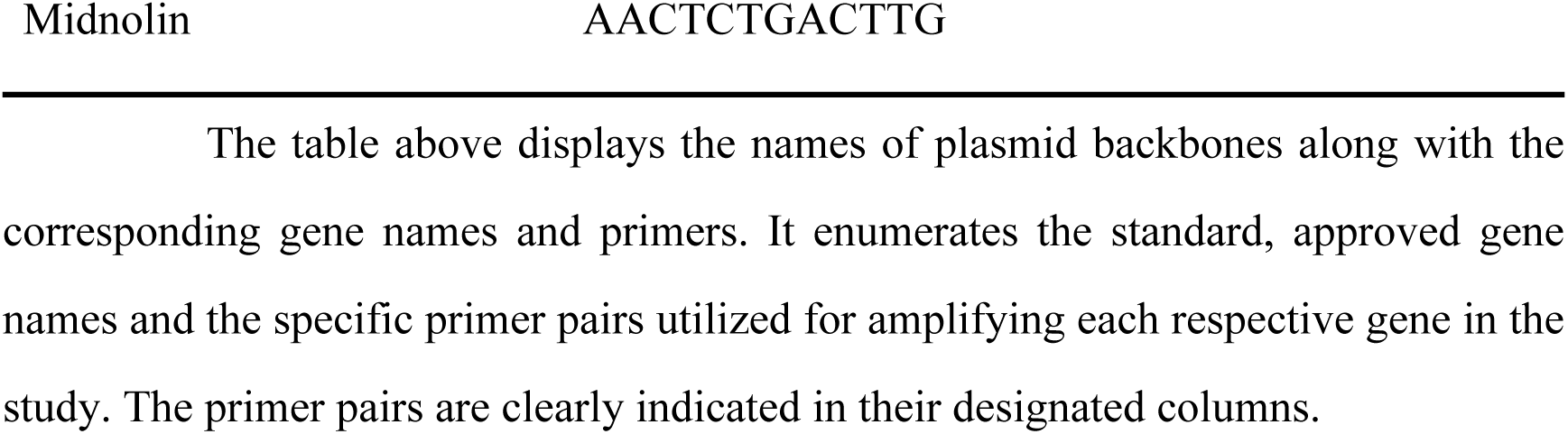
Gene Names and Corresponding Primer Pairs.

